# Unbiased CRISPR Synthetic Lethal Screening for Genetic Vulnerabilities in Succinate Dehydrogenase (SDH)-loss Model of Paraganglioma

**DOI:** 10.1101/2025.08.26.672429

**Authors:** Fatimah J. Al Khazal, Michael J. Emch, Cristina M. de Araujo Correia, Judith Favier, John R. Hawse, L. James Maher

## Abstract

Succinate dehydrogenase (SDH)-deficient paraganglioma and pheochromocytoma (PPGL) are rare neuroendocrine tumors for which no effective targeted therapies currently exist. To uncover new potential therapeutic targets, we performed an unbiased CRISPR-Cas9 genetic screen in immortalized mouse chromaffin cells (imCCs) with and without *Sdhb* loss. Our screen identified genes that differentially affect cell proliferation in *Sdhb*-deficient versus normal imCCs. Notably, several subunits of the transcriptional Mediator complex emerged as potential tumor suppressors, as their loss selectively promoted growth of *Sdhb*-deficient cells. *Most strikingly*, we found that the neddylation pathway—required for ubiquitin-mediated selective protein degradation—plays a critical role in controlling cell growth and survival in *Sdhb*-deficient imCCs. Specifically, loss of the neddylation regulator *Ube2m* led to increased proliferation, while loss of *Ube2f* suppressed growth of *Sdhb*-deficient imCCs. Consequently, global neddylation inhibitor MLN4924 (Pevonedistat) and UBE2F-CRL5 axis inhibitor HA-9104 were shown to downregulate neddylation, suppressing UBE2F activity and selectively inhibiting growth of *Sdhb*-deficient imCCs. This unexpected result highlights the neddylation pathway as a promising druggable vulnerability in this cell culture model of SDH-deficient PPGL.

## Introduction

Pheochromocytomas (PHEO) and paragangliomas (PGL) are rare neuroendocrine tumors that arise from paraganglia, cell clusters of embryonic neuroectodermal origin that can secrete catecholamines and other neurotransmitters [1]. “PHEO” designates PGL tumors that develop from chromaffin cells in the adrenal medulla, whereas “PGL” is most often is applied to extra-adrenal tumors of the same origin [2]. Together PHEO and PGL are here termed “PPGL.” Intriguingly, a clinically important PPGL subtype, both molecularly and morphologically, is driven by deficiency in succinate dehydrogenase (SDH) [3]. SDH-deficient PGL tumors arise in asymptomatic heterozygous *Sdh* variant carriers through loss of heterozygosity (LOH) of any of the four genes encoding the four subunits of SDH (SDHA, SDHB, SDHC, or SDHD) or assembly factor (SDHAF2). Strikingly, the immunohistochemical signal for the SDHB subunit typically becomes negative following the loss of any SDH subunit, making SDHB loss a relatively reliable histological marker of SDH-loss PPGL [4, 5]. SDHB-deficient tumors are notable in their clinical aggressiveness, propensity to metastasis and recurrence, and higher morbidity and mortality compared to other forms of hereditary and sporadic PPGLs [6].

Conventional treatment for PPGL involves surgical resection, which can be challenging depending on tumor localization, vascularity, and potential for metastasis [7]. For surgically inaccessible tumors, a combination of radiation therapy (external beam or radioisotope peptide receptor radionuclide therapy [PRRT]), ablation, or embolization may be used with or without chemotherapy and targeted drugs to control excess blood catecholamines [8]. A major gap in PPGL research lies in developing treatments targeted to molecular pathways and genetic vulnerabilities specific to PPGL tumor sub-types, decreasing the need for invasive medical interventions [9]. One targeted therapeutic strategy is treatment with HIF-2α inhibitor Belzutifan. This recently-approved drug treats PPGL tumors with a distinct pseudohypoxic signature driven by HIF-2α overexpression secondary to HIF prolylhydroxylase inhibition by succinate accumulation in SDH-deficient PPGL, or by VHL ubiquitin ligase loss in VHL-deficient PPGL [10]. There have been additional limited attempts to exploit the unique genetic and epigenetic vulnerabilities of SDH-deficient PPGL tumors [10, 11]. It remains to be seen whether Belzutifan will have long-term positive therapeutic impact.

Here we deploy an unbiased genome-wide CRISPR-Cas9 synthetic lethal genetic screen in a mouse adrenal medulla-derived cell model of *Sdhb*-loss and its wild-type counterpart to uncover genetic vulnerabilities in the form of genes that, once deleted, selectively reduce survival of *Sdhb*^-/-^ cells but not *Sdhb*^+/+^ cells. In the course of this work, genes whose loss selectively enhance growth of *Sdhb*^-/-^ cells are also revealed. To our fascination, we uncovered complementary impacts of loss of two enzymes, UBE2M and UBE2F, that play roles in the NEDD8 conjugation (neddylation) pathway required for ubiquitin-mediated targeted protein degradation. Eliminating their respective genes induced opposite effects: selective synthetic survival or lethality in SDHB-loss cells, respectively, while SDH-expressing cells tolerated both deletions. These results were further validated using shRNA expression to silence these targets, with similar results. Finally, we show that *Sdhb*^-/-^ imCCs, but not SDH-expressing cells, are selectively sensitive to two neddylation inhibitors, MLN4924 (Pevonedistat) and HA-9104, believed to reduce UBE2F-CRL5 levels. These results point to the unexpected potential of the neddylation pathway as a druggable target in *Sdhb*-deficient cells, motivating further characterization and testing in other models of SDH-loss PPGL.

## Results

### Unbiased CRISPR screen identifies both synthetic lethal and synthetic growth genes that selectively affect SDH-loss imCCs

To identify genes whose loss is synthetically lethal with *Sdhb* loss in an adrenal medulla-derived immortalized murine cell line, but not in corresponding SDH-expressing cells [12–14], we performed a genome-wide CRISPR-Cas9 knockout screen using a mouse CRISPR knockout pooled library (Brie) [15], which carries 78,637 single guide RNAS (sgRNAs) targeting 19,674 genes (∼4 sgRNAs per targeted gene). The library also contains an additional 1000 “dummy” guides that serve as controls. To employ this approach, we introduced the sgRNA library by lentiviral transduction into Cas9-expressing imCCs (Supplemental Fig. S1A) and applied appropriate antibiotic selection and cell counting to ensure 600-fold coverage of the library at a MOI of ∼0.4, so that only sgRNA-expressing cells survive in the experimental pool for each line. Under these conditions, about 6% of cells will express two or more sgRNAs. The low frequency of such events, coupled with the extreme rarity that more than one cell would be transduced by the same two guides makes the condition acceptable for screen interpretation. Each transduced cell pool (SDH-loss and SDH-expressing) was then cultured for ten cell doublings with half the cell population collected at each doubling. Genomic DNA was extracted from all 20 samples, and PCR was used to amplify bar-coded sgRNA sequences from the genomic DNA. Deep sequencing of the resulting pooled DNA was then performed on an Illumina NovaSeq 6000 S4 flow cell, and the resulting data were sorted via the caRpool R package for sequence alignment against Brie CP0044 and then analyzed using the MAGeCK algorithm to identify genes whose sgRNAs were consistently either depleted (synthetic lethality) or enriched (synthetic growth) in the final cell population. This experimental workflow is outlined in Fig. 1A. Leading gene candidates from the screen are ranked based on the combined effects of their associated sgRNAs (Fig. 1B and 1C), specifying that identified sgRNAs are collectively ranked higher than what would be expected by random chance. Genes whose deletion adversely affected cell proliferation (synthetic lethality) are shown in Fig. 1B. Among these candidates, NEDD8 conjugating enzyme *Ube2f* in the neddylation pathway related to ubiquitin-mediated targeted protein degradation stood out as the most significant hit. This candidate also immediately caught our attention because its upstream regulatory antagonist in the same neddylation pathway, *Ube2m* [16], appeared as one of the top synthetic growth targets (Fig. 1C). Thus, loss of *Ube2f* expression was selectively lethal to SDH-deficient cells (not SDH-expressing cells), while loss of *Ube2m* expression selectively enhanced growth of SDH-deficient cells (not SDH-expressing cells).

**Figure 1.**
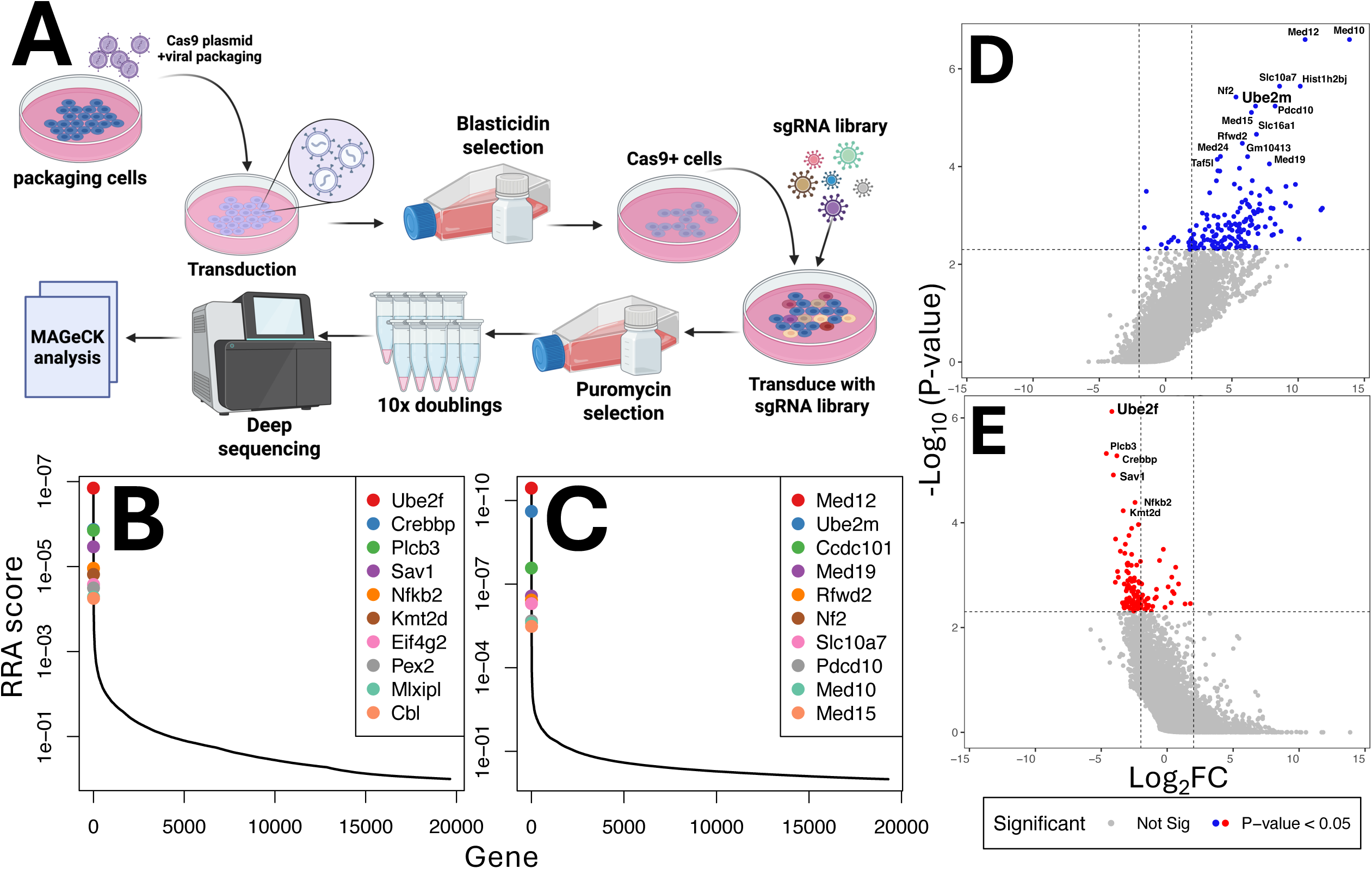
Genome-wide CRISPR/Cas9 knockout screen to identify genes that modulate proliferation in SDHB-loss vs. SDH-expressing immortalized mouse chromaffin cell lines. **(A)** Schematic workflow of the pooled CRISPR/Cas9 gene knockout screen. Cas9-expressing chromaffin cells were generated via lentiviral transduction and blasticidin selection. Cells were then transduced with a genome-wide sgRNA library, selected with puromycin, and cultured for 10 population doublings before genomic DNA extraction. sgRNA abundance was quantified at initial and final times by deep sequencing, and data were analyzed using the MAGeCK pipeline. **(B)** Ranked Robust Rank Aggregation (RRA) scores of the most significantly depleted genes from the screen. **(C)** Ranked RRA scores of the most significantly enriched genes from the screen. **(D)** Volcano plot of CRISPR screen results showing significantly depleted genes (blue) versus non-significant genes (gray). The x-axis represents log₂(fold change) (Log₂FC) in sgRNA abundance, and the y-axis shows –log₁₀ *P* values from MAGeCK analysis. **(E)** Volcano plot showing significantly enriched genes (red) versus non-significant genes (gray).

Further evidence that targeted protein degradation can be a unique dependency of SDH-loss cells is the observation that *Ntan1* appears among top 50 synthetic lethal genes in SDH-loss imCCs (deposition pending). *Ntan1* encodes an N-terminal asparagine amidase required within the N-end rule pathway of targeted protein degradation, converting stabilizing N-terminal asparagine residues to aspartic acid, a substrate for conjugation to a destabilizing N-terminal arginine residue, facilitating ubiquitylation and proteasomal degradation [17].

Intriguingly, selective synthetic growth genes included several encoding subunits of the transcriptional Mediator complex (Med12, 10, 15 and 19), a large, multi-subunit protein complex that plays a crucial rule in transcriptional regulation [18]. This result suggests that activation of certain genes that are tumor suppressors in the context of SDH loss are particularly dependent on certain Mediator subunits. To complement RRA score ranking, which reflects combined effects of gene-specific effects of sgRNAs, we also calculated p-values to assess the statistical significance of these effects (Fig. 1D-E). Together, these results confirm unanticipated synthetic lethal and synthetic growth effects of neddylation pathway E2 regulator gene deletion in the context of *Sdhb* loss. While other listed candidate genes are also intriguing and merit future study, the presence of complementary neddylation enzymes among top candidates in the synthetic lethal and synthetic growth gene lists drew our attention to prioritize analysis of these genes, particularly synthetic lethal candidate *Ube2f*.

### Validation of neddylation screen hits by targeted shRNA knockdown

To validate the top synthetic lethal target (Ube2f) and its complementary antagonistic partner identified in the screen as synthetic growth target (Ube2M), we performed an independent shRNA knockdown validation of these E2 family enzymes to test if the synthetic effects could be reproduced by mRNA knockdown [19]. Target proteins persisted in both experimental lines regardless of *Sdhb* status after commercial siRNA treatment, so this was deemed an inadequate approach [20]. Using alternative expression of shRNA constructs targeting *Ube2m* and *Ube2f* mRNAs, we were able to create stable lines displaying detectable knockdown of these proteins (Fig. 2D, lanes 2,5 and 3,6, respectively). Interestingly, robust knockdown required ∼8-10 cell doublings after transduction (Supplemental Fig. S2 and Fig. 2D). At that point, shRNA targeting of *Ube2m* and *Ube2f* significantly and selectively affected growth of *Sdhb^-/-^*, but not SDH-expressing imCCs (Fig. 2A-B). The results are shown as effects on absolute growth rate (Fig. 2A) and normalized to the effect of non-targeted shRNA (shNT; Fig. 2B) to account for the much slower growth rate of SDH-loss imCCs [13]. After >8 doublings, the *Ube2f* stable knockdown line did not show profound apoptosis presumably because any immediate synthetic lethality that occurred before bulk *Ube2f* knockdown could be detected by western blotting resulted in early loss of that cell subset. *Ube2f* knockdown cells capable of persisting until UBE2f protein levels were detectably reduced in western blots presumably had adapted to tolerate UBE2F loss with reduced growth rate but not frank apoptosis.

**Figure 2.**
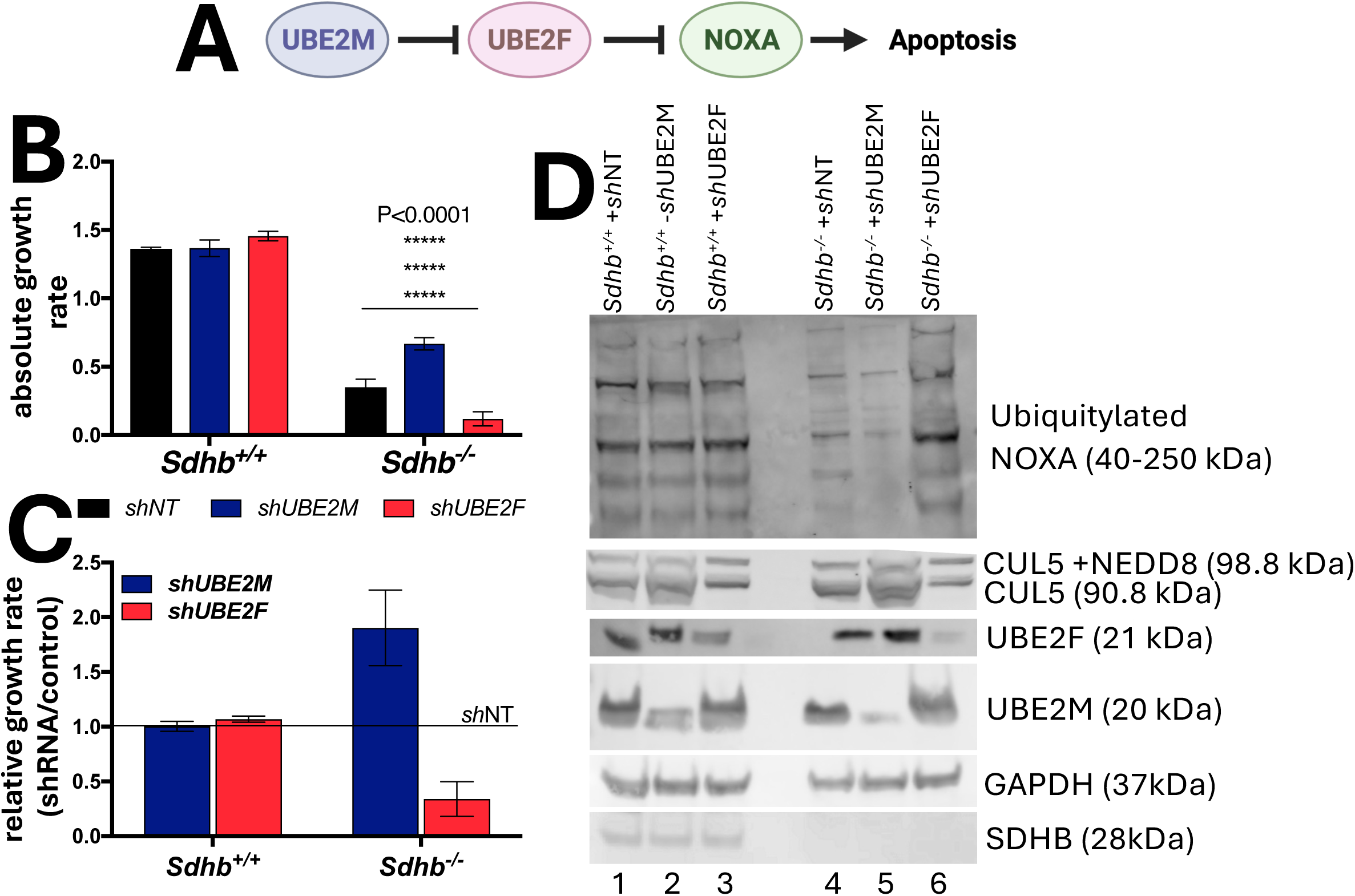
shRNA-based validation of differential effects of UBE2M and UBE2F knockdown on proliferation of Sdhb^+/+^ and Sdhb^-/-^ imCCs. **(A)** Schematic illustration of the regulatory relationship between UBE2M and UBE2F in regulating apoptosis. **(B)** Absolute growth rates of *Sdhb^+/+^* and *Sdhb^-/-^* imCCs following knockdown with non-targeting control (shNT), shUBE2M, or shUBE2F. UBE2M knockdown selectively enhances growth in *Sdhb^-/-^* cells, while UBE2F knockdown strongly suppresses growth in both genotypes. P-values are calculated using Bonferroni correction for multiple t-tests and the number of asterisks indicates degree of significance. **(C)** Relative growth rates from (B) normalized to shNT controls, highlighting the lineage-specific increase in proliferation upon UBE2M knockdown in *Sdhb^-/-^* cells. Error bars account for propagation of error. **(D)** Western blot analysis of ubiquitylated NOXA (high-molecular-weight smear, ∼40–250 kDa), CUL5, UBE2F, UBE2M, GAPDH (loading control), and SDHB in *Sdhb^+/+^* and *Sdhb^-/-^* imCCs after knockdown with shNT, shUBE2M, or shUBE2F. Protein depletion was observed only 8-10 cell doublings after treatment. The pattern suggests altered CUL5-associated substrate ubiquitylation upon UBE2F loss and stabilization of UBE2F substrates upon UBE2M depletion in the *Sdhb^-/-^* background.

To further analyze these results, western blotting for apoptotic factors believed to be the targets of the *Ube2m*/*Ube2f* neddylation pathway was undertaken. *Ube2f* knockdown cells were characterized by the absence of the pro-apoptotic protein NOXA at its native molecular weight (6-15 kDa), though increased expression of ubiquitylated forms of NOXA in response to lower levels of UBE2F (Fig. 2D, lane 6) may still explain why these *Sdhb^-/-^* cells grew poorly (Fig. 2B-C) [21]. CUL5, a scaffold protein that interacts with UBE2F and other proteins to form the CUL5 RING ubiquitin ligase 5 (CRL5) complex, was also observed to be reduced upon knockdown of UBE2F expression (Fig. 2D, lanes 3 and 6; [22, 23]). We note that our CRISPR screen also identified CUL5 among the top 20 synthetic lethal targets (deposition pending), strikingly suggesting that the SDHB-loss genetic vulnerability in question involves a client of CRL5 (containing UBE2F-CUL5), possibly the pro-apoptotic protein NOXA [24].

### Pharmaceutical targeting of the neddylation pathway selectively impairs growth of SDH-deficient imCCs

To further provide evidence that the neddylation pathway of targeted protein degradation represents a selective conditional dependency in *Sdhb* loss imCCs, we tested two commercially available neddylation inhibitors. MLN4924 blocks neddylation globally by inhibiting NEDD8-Activating Enzyme (NAE) to prevent NEDD8 activation. This prevents neddylation of any protein targets. We also tested selective inhibitor HA-9104, which blocks the V30 pocket of UBE2F to prevent binding of CUL5 (Fig. 3A) [22, 25]. Comparing the two agents, treatment with HA-9104 resulted in less toxicity to control SDH-expressing imCCs, while *Sdhb^-/-^* imCCs were significantly more sensitive to inhibition. Peak selective inhibition of SDH-loss imCCs was observed between 2.5 and 5 µM HA-9104 (Fig. 3B). Under these conditions, western blot analysis of key neddylation components shows striking loss of UBE2F and CUL5 starting at 12-24 h post-treatment (Fig. 3B and Fig. 3D, lanes 3,4,7,8). Similarly, *Sdhb^-/-^* imCCs were considerably less tolerant of MLN4924 treatment at all tested concentrations higher than 1 µM, with SDH-expressing cells demonstrating the least toxicity at 1.5 µM (Fig. 3C). A parallel western blot analysis supports loss of global neddylation through progressive and direct degradation of NEDD8 rather than E2 enzymes (Fig. 3E, lanes 3,4,7,8). Notably, levels of the pro-apoptotic regulator NOXA are selectively elevated by neddylation inhibitors only in SDH-loss cells (Fig. 3F, lane 8 in left and right panels). Together, these results demonstrate that pharmaceutical inhibition of neddylation, both globally and through targeted UBE2F inhibition, selectively inhibits growth of SDH-loss imCCs.

**Figure 3.**
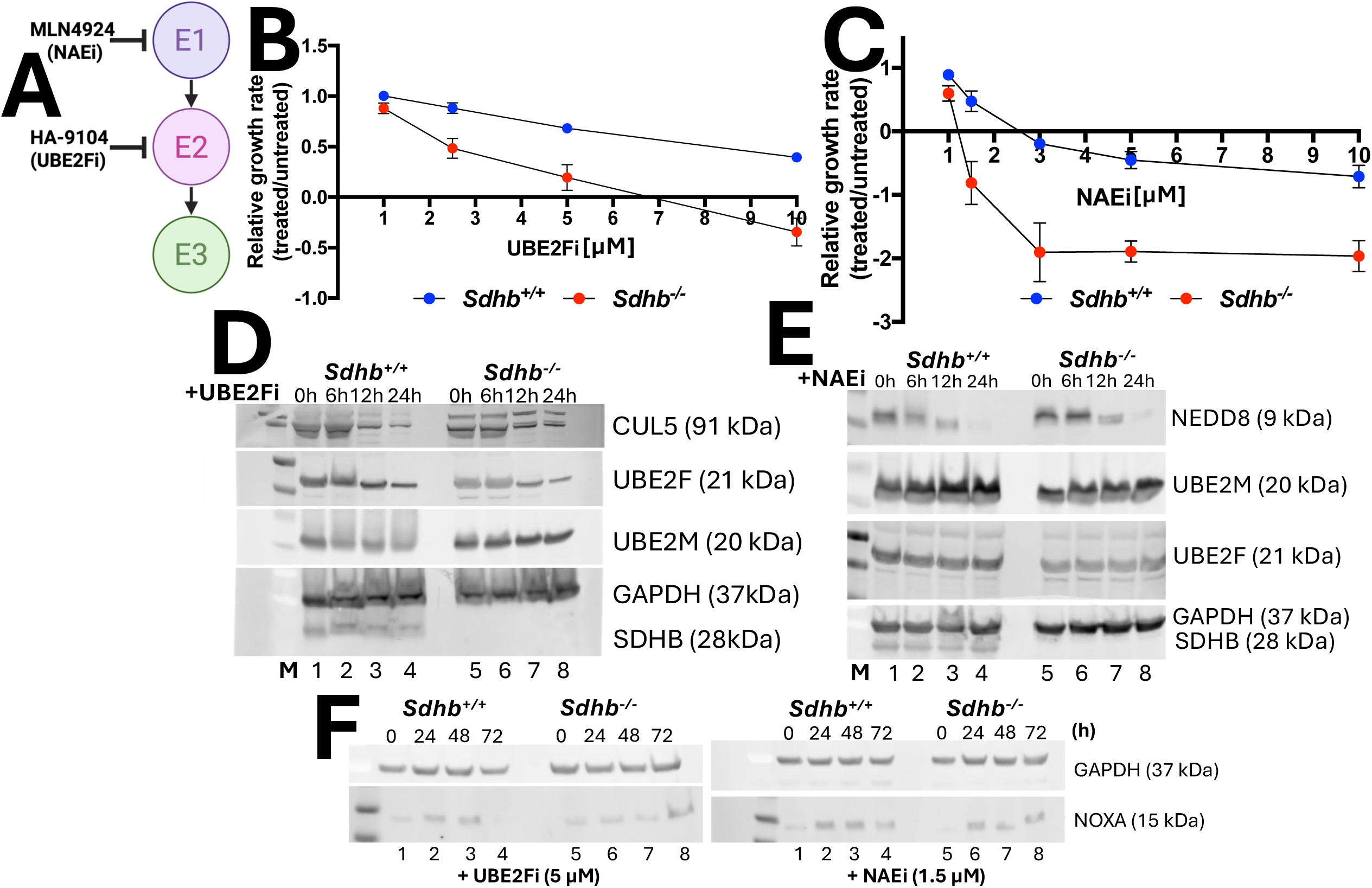
Pharmacological inhibition of UBE2F and the neddylation pathway in Sdhb^+/+^ and Sdhb^-/-^ cells yields selective growth inhibition of Sdhb^-/-^ imCCs. **(A)** Schematic figure of the neddylation cascade and pharmacological targeting at the E1 (NAE1) or E2 (UBE2F) levels. Drugs were used to block UBE2F (UBE2Fi) or NEDD8-activating enzyme (NAEi). **(B)** Dose–response data showing relative growth rates (treated/untreated) of *Sdhb^+/+^ and Sdhb^-/-^* cells following treatment with the UBE2F inhibitor (UBE2Fi) for 2 doublings. **(C)** Immunoblot analysis of CUL5, UBE2F, UBE2M, GAPDH, and SDHB after UBE2Fi treatment for 0, 6, 12, and 24 h, showing progressive reduction in CUL5 neddylation and changes in UBE2F protein levels. **(D)** Dose– response data showing relative growth rates after treatment with the NEDD8-activating enzyme inhibitor (NAEi) for 72 h. Both *Sdhb^+/+^ and Sdhb^-/-^* cells exhibit growth suppression, with *Sdhb^-/-^* cells showing a stronger effect. **(E)** Immunoblot analysis of NEDD8, UBE2M, UBE2F, GAPDH, and SDHB after MLN4924 treatment for 0, 6, 12, and 24 h, confirming global inhibition of neddylation through suppression of NEDD8 and no changes in E2 enzyme levels.

### DepMap analysis reveals pan-dependencies on the neddylation pathway in Sdhb-loss lines

By comparing our screen results to the Dependency Map (DepMap) essentiality scores from the Broad Institute analysis of relevant tumors with SDH loss [26], we noted both tissue-specific and *Sdhb* loss-specific dependencies. This analysis validated the crucial nature of certain genes and pathways, while also providing insight into the selective pressures of SDH loss unique to different cancer types. Fig. 4 provides robust evidence for the essentiality of key neddylation pathway components in three different tumor types with SDH loss: Kidney Renal Clear Cell Carcinoma, Lung Adenocarcinoma, and Stomach Adenocarcinoma. Interestingly, the DepMap scores for UBE2M, UBE2F, and CUL5 consistently fall in the negative range across all three cancer models (Fig 4A-C). This finding strongly suggests that the neddylation pathway is a fundamental and non-redundant process required for the survival and proliferation of these diverse cancer cells. This cross-cancer validation reinforces the neddylation pathway as a promising and broadly applicable therapeutic target. Conversely, these data also highlight genes that are not essential for growth of these cancers. The pro-apoptotic protein NOXA shows a positive DepMap score in all three cancer types, indicating that its depletion has minimal impact on cell growth in these tumors. This suggests that these malignancies are not reliant on suppression of NOXA for survival and they may possess alternative mechanisms to evade apoptosis. Interestingly, mediator protein complex subunits MED10, MED12, MED15 and MED19 are found to be synthetic lethal genes when deleted in the absence of SDH in the tested tumors, as indicated by negative DepMap scores, suggesting essential roles in the viability of the tested cancers. The fact that we found loss of these Mediator subunits to cause selective *growth* of SDH-loss imCCs, and that elevating NOXA levels to be inhibitory to SDH-loss imCCs points to the cell-type dependence of these genes [27]. The variation in the exact sensitivity scores and the ranking of other genes across the three cancers further demonstrates the importance of specific tumor context when considering targeted therapies.

**Figure 4:**
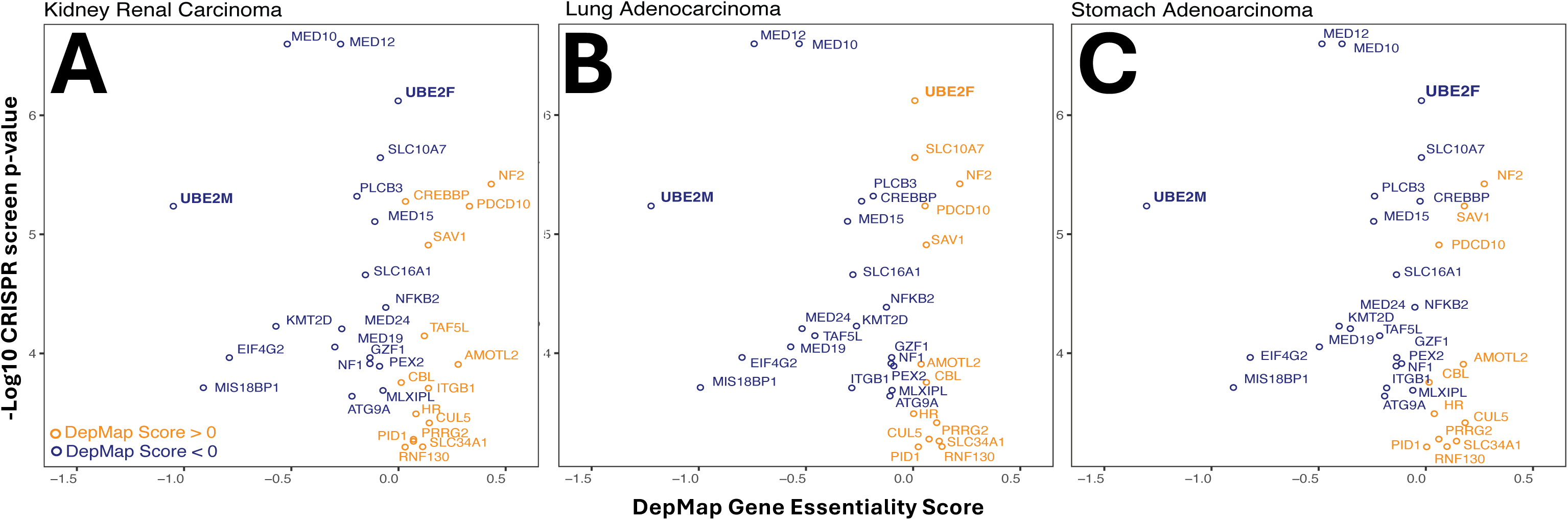
Comparison of DepMap essentiality scores across various Sdhb-deficient cancer lines. Scatter plot depicts the p-value (log_10_) for top differential genes detected from the SDHB screen and the DepMap gene essentiality scores (chronos scores) in kidney renal cell carcinoma (A), lung adenocarcinoma (B) or stomach adenocarcinoma (C) (n=57). Genes are colored based on their average essentiality score, orange genes core > 0; orange and, genes core <0; blue. The Spearman rho coefficient= 0.250 and p-value=0.160.

## Discussion

The investigation reported here combines an unbiased genome-wide CRISPR-Cas9 screen to identify growth dependencies unique to SDH-loss cells, followed by targeted validation using shRNA knockdown and pharmaceutical inhibition. The results provide compelling evidence that the neddylation pathway of targeted protein degradation emerges as an important dependency in this cell culture model of SDH-deficient PPGL. This finding was unexpected to us, as we anticipated gene candidates related to metabolism. The result has critical implications for understanding metabolic and signaling adaptations in SDH-loss cells, and for developing novel therapies.

Neddylation is a post-translational modification in which an E1 activating enzyme (NAE) activates the small ubiquitin-like NEDD8 protein, allowing its conjugation by E2 conjugation enzymes to E3 cullin and non-cullin protein substrates, facilitating downstream ubiquitin-dependent targeted protein degradation [25]. By recruiting E3 ligases and targeting cullin and non-cullin targets, E2 NEDD8 conjugating enzymes UBE2M and UBE2F are upstream regulators of various crucial biological processes including cell cycle arrest, DNA damage response, hypoxia and stress responses and tumorigenesis that depend on targeted protein degradation [24, 25, 28–30].

In our specific model of *Sdhb*-loss adrenal medulla-derived murine cells, it appears that the neddylation pathway is a double-edged sword. Loss or inhibition of UBE2F activity selectively impairs growth of *Sdhb*-deficient cells, whereas loss or inhibition of its antagonizing E2 enzyme and negative regulator, UBE2M, selectively promotes the growth of SDH-loss imCCs (Fig. 1B-E and Fig.2). Strikingly, while both enzymes have previously been discussed as synthetic dependencies in various malignancies, they are both often described as synthetic lethal targets whose loss is associated with slower tumor progression, higher sensitivity to chemical targeting or cell cycle arrest [24, 30–34]. This is in contrast to our results in SDH-loss imCCs, in which *Ube2m* loss selectively promotes growth of SDH-loss imCC, presenting an intriguing context-specific dependency (Fig. 4). Thus, while loss of both UBE2M and UBE2F have been individually implicated as synthetic lethal vulnerabilities in some cancers, their identification as demonstrating opposing synthetic effects is novel. This result clearly supports the model of Zhou et al. shown in Fig. 2A wherein UBE2F targets the destruction of pro-death NOXA, whose low levels are selectively important for survival of SDH-loss cells, and UBE2M participates in the targeted destruction of UBE2F in these cells [16]. Thus, SDHB-loss imCCs are inherently intolerant of *Ube2f* loss. In contrast, when *Ube2M* is lost, *Ube2F* is preserved as an inhibitor of a pro-death factor, likely NOXA, to which SDH-loss cells are highly sensitive.

The role of *Ube2m* as a tumor suppressor gene has previously been linked to its function as a p53 negative regulator [32, 35]. Interestingly, the *Sdhb*-loss imCCs express 10-fold less p53 that SDH-expressing imCCs, possibly explaining their behavior upon loss of *Ube2M* [36]. Our *Sdhb*-loss imCC model is also characterized by a severe pre-existing metabolic defect, slow growth, extensive chromosomal and DNA copy number abnormalities, pseudohypoxia-related stress and profound ATP-deficiency [13, 36]. These widespread defects may lead to a unique responses relative to other cell types. The observed growth advantage upon UBE2M loss likely arises from loss of its negative regulatory control over UBE2F, enhancing cell proliferation as seen in Fig. 2B-C, presumably by permitting UBE2F to degrade one or more cell death proteins.

Our screen shows that loss of neddylation E2 enzyme UBE2F is synthetically lethal with SDH loss in imCCs (Fig. 1B, Fig. 3B-E). Fortunately, pharmaceutical agents targeting the neddylation pathway at the E1 and E2 levels are available [22, 37], and improved inhibitors are being developed. MLN4924 (NAEi), a global neddylation inhibitor currently under investigation in multiple phase I clinical studies on several malignancies. This agent has been shown to exert anticancer activities by inducing cell cycle arrest, apoptosis, senescence and autophagy in a cell-type and context dependent manner [28, 37, 38]. MLN4924 inhibits the E1 enzyme NAE1, which is the first and most critical step in the entire neddylation cascade, effectively downregulating global neddylation activity, leading to the inappropriate accumulation of sensitive protein substrates and ultimately causing cell death. [28, 37]. It is therefore not surprising that MLN4924 has baseline toxic effects regardless of *Sdhb* status, though *Sdhb*-loss cells remain more sensitive to MLN4924 treatment, proving that global targeting of neddylation is selectively more harmful in the context of cellular stresses due to *Sdhb* loss (Fig. 3D-E). More notably, targeted chemical inhibition of UBE2F by HA-9104 [30], shows significantly less toxicity in SDH-expressing control imCCs compared to SDH-loss cells, further supporting the notion that UBE2F-related functions become dependencies upon *Sdhb*-loss (Fig. 3B-C).

Though UBE2M is well characterized in the literature, less is known of UBE2F functions and E3 regulatory clients. In the context of this work, future identification of specific substrates of UBE2F will be crucial to understanding why SDH loss sensitizes cells to the loss of this E2 enzyme. For example, further research might explore possible links between the pseudohypoxia and metabolic stress caused by SDHB loss and the specific regulatory role of neddylation.

Based on the data shown here, we hypothesize that NOXA, a pro-apoptotic protein whose degradation is controlled through UBE2F and SDH-loss cells are hypersensitive to NOXA levels levels [16]. We show that NOXA levels fluctuate significantly in response to changes in UBE2F levels (Fig. 2D and Fig. 3F). NOXA also tends to persist longer in *Sdhb^-/-^* vs. *Sdhb^+/+^* cells treated with neddylation inhibitors for 72 h compared (Fig. 3F lanes 4 and 8). We show that stable shRNA knockdown of UBE2F in SDH-loss cells can result in tolerance to UBE2F loss over multiple cell doublings, but growth rate is severely attenuated and NOXA levels (mainly ubiquitylated forms) increase. Pharmaceutical inhibition of UBE2F or NAE1 results in robust, sudden loss of neddylation activity indicated by loss of NEDD8 and UBE2F/CUL5 (Fig. 3B and 3C), accompanied by parallel increased expression in NOXA at its anticipated molecular mass (Fig. 3F, lane 8 in left and right panels). This suggests that NOXA-driven apoptosis is the plausible mechanism by which UBE2F loss is synthetically lethal with SDH loss.

The implication of this work is that SDH-loss imCCs are exquisitely sensitive to apoptosis driven by NOXA levels while SDH-expressing imCCs are not. What might account for this? We propose that the profound ATP deficiency of SDH-loss cells is a primary consideration [36]. In this model, supported by data from ATP-deficient IDH-mutant tumors [39], diminished ATP production in SDH-deficient imCC leads to diminished mTOR pathway signaling, and lower levels of NOXA antagonist MCL1, thus diminishing the ability of cells to tolerate NOXA upregulation, and enhanced sensitivity to inhibitors of pathways parallel to the MCL1 pathway. Future studies will be necessary to pursue these possibilities.

An additional possible follow-up study would involve a second genome-wide CRISPR screen on SDHB-loss and SDHB-expressing imCCs under physiologically-relevant oxygen and nutrient conditions. This would allow us to confirm the observed synthetic lethality and determine if it persists in relevant lower physiological oxygen and human plasma-like nutrient levels [21, 24].

## Materials and Methods

### Cell line maintenance

SDH-expressing and SDH-loss imCCs have previously been described and characterized for doubling time and optimal culture conditions, which were sustained following the creation of Cas9-expressing lines [13]. Throughout the processes of establishing additional stable cell lines, establishing appropriate kill curves for puromycin and blasticidin and the CRISPR synthetic lethal screening process, culture conditions were: 37 °C, 95% humidity in room air (21% O_2_) with 5% CO_2_. Growth media consisted of high glucose DMEM containing GlutaMAX™ (Gibco #10566016), 10% heat-inactivated FBS (Gibco #10082147) and a 0.5 mg/mL final concentration of pen/strep (Gibco #15140122). Media were further supplemented with nonessential amino acids [100 μM final concentration each of glycine, alanine, asparagine, aspartic acid, glutamic acid, proline, and serine (Gibco #11140035)], 1 mM sodium pyruvate (Gibco #11140035) and 10 mM HEPES buffer (Gibco #15630130). For cell line expansion, cells were supplied with fresh media every other day and replated based on their doubling rate when 80-90% confluence was reached. During the screen, each line was split in half after each doubling (24 h for *Sdhb^+/+^*cells; 96 h for *Sdhb^-/-^* cells) and half the cells were collected for DNA extraction.

### Generation of polyclonal Cas9-expressing imCC lines

Parental imCC lines were made to express Cas9 by transduction with 3^rd^ generation lentiviral particles carrying codon-optimized *S. pyogenes* Cas9 protein and blasticidin resistance driven from the EFS promoter (Addgene # 52962-LV). Briefly, SDHB-expressing and SDHB-loss cells were seeded into 6-well plates at 50% confluence (50,000 cells) and allowed to adhere overnight. The next day, growth medium was removed, and the cells were incubated in polybrene-containing media (10 µg/mL final concentration) for 2 h prior to viral transduction. Three h later the wells were treated with lentivirus at an MOI of 0.4, 0.8, 1, or 2, leaving 2 wells as counting and blasticidin controls for each line. The following day (18 h after transduction), medium was aspirated, wells were washed thrice with PBS before adding selection media containing 10 µg/mL blasticidin [13]. Wells demonstrating 40-50% cell survival (compared to counting control) following 7 d of selection (at which point blasticidin controls had reached 100% cell death) were expanded and frozen. This approach deliberately favors polyclonal stable cell populations to avoid any peculiartities that might be associated with a clonal population. Western blotting was used to confirm Cas9 expression compared to parental lines (Supplemental Fig. S1).

### CRISPR-Cas9 screen

The synthetic lethal screen was carried out using Mouse CRISPR Knockout Pooled Library (Brie) lentiviral prep (Addgene #73633-LV) at 600-fold coverage and a MOI of 0.4 [15]. SDH-loss cells have previously been characterized for their relative insensitivity to viral transduction. As such, the equivalent functional MOI in SDH-loss cells was found to require 1.8-fold higher levels of viral particles to achieve the same rate of infection as for SDH-expressing cells. For this process, ∼18-h incubation with 10 µg/mL of DEAE-dextran containing media prior to infection with Brie library was required. The following day, cells were transduced with the appropriate volume of viral particles in DEAE-dextran containing media and left to incubate for 18-20 h. The next day, viral particles-containing media were removed and safely discarded, cells were washed three times in PBS and provided normal growth media to allow for recovery and expression of the puromycin resistance marker. At 48 h post-transduction cells were re-plated to prevent clustering, and growth medium was spiked with the appropriate concentration of Puromycin Dihydrochloride (Gibco #A1113803) for SDH-expressing and SDH-loss cells (2.5 and 5 μg/mL, respectively) [40]. Fresh selection media was supplied daily, and a puromycin control flask (non-transduced cells) was used to mark the appropriate selection endpoint at 72 h post-treatment, at which point half the cells were collected as T_1_ (first doubling) samples, while the remaining cells from each line were replated into fresh flasks. Control cells were split in half and collected every 24 h while SDH-loss cells were collected every 96 h, corresponding to their respective doubling rates. Collected cells were washed with PBS and frozen at -20 °C until all samples from all ten doublings had been collected.

### DNA extraction and PCR-facilitated barcoding

An Illumina next-generation sequencing protocol was followed for nucleic acid isolation, followed by library preparation. DNA extraction and purification was performed using DNeasy Blood & Tissue Kit (Qiagen #69504) according to manufacturer protocol.

For library amplification, a cocktail of 8 Illumina staggered P5 primer (5’-A_2_TGATACG_2_CGAC_2_AC_2_GAGATCTACACTCT_3_C_3_TACACGACGCTCT_2_C_2_GATCT[N]T_2_GT G_2_A_3_G_2_ACGA_3_CAC_2_G-3’) and one unique indexing P7 primer was used per sample (5’-CA_2_GCAGA_2_GACG_2_CATACGAGAT*NNNNNNNN*GTGACTG_2_AGT_2_CAGACGTGTGCTCTT C_2_GATCT_2_CTACTAT_2_CT_3_C_4_TGCACTGT-3’). The following machine cycle steps were followed: denaturation: 95 °C for 1 min (initial) then 95 °C for 30 s; annealing: 53°C for 30 s; extension: 72°C for 30 s; repeat: Steps 2-4 for 34 cycles; final Extension: 72°C for 10 min. Standard agarose gel electrophoresis (2%) was used to check the size and quality of amplicons across samples (expecting a product size of 354 bp). AMPure XP beads (Beckman Coulter #A63880) were used according to manufacturer protocol to clean up and concentrate the amplicon samples prior to submission.

### Next Generation Sequencing (NGS)

Sample submission was made to the Mayo Clinic Division of Laboratory Genetics and Genomics, sequencing facility where libraries were sequenced on an Illumina NovaSeq 6000 S4 flow cell, providing ∼20 billion total reads.

### Generation of shRNA stable imCC lines

MISSION shRNA Lentiviral particle system reagents(Sigma Aldrich #SHCLNV) were ordered against *Ube2m* (clone ID #TRCN0000317874) and *Ube2f* (clone ID #TRCN0000098574) and used for stable protein knockdown of parental imCC lines, coupled with pLKO.1-puro non-mammalian shRNA as transduction control (Sigma Aldrich #SHC002VN). Transduction and selection were performed on *Sdhb^+/+^* and *Sdhb^-/-^* imCCs according to the manufacturer’s protocol with the appropriate puromycin concentration (2.5 and 5 μg/mL, respectively) for each line for 3 d. Transduced lines were monitored for protein reduction via western blotting. Significant target protein reduction was confirmed only after 8-10 doublings for both lines, suggesting high protein stability even after mRNA knockdown.

### Pharmaceutical validation using neddylation inhibitors

Pevonedistat (MLN4924) and HY-160799 were obtained from MedChemExpress in the form of 1 mL aliquots of 10 mM drug in DMSO (ID #HY-70062 and HY-160799). Initial pilot studies for each drug were conducted to determine the appropriate concentration range and treatment protocol. Regardless of SDHB status, both imCC lines exhibited resistance to MLN4924 at previously reported concentrations between 0.1-1 µM for up to 7 d– but were responsive within the 1-10 µM range (Supplemental Fig. S3 C-D) [28]. Both lines showed relatively consistent sensitivity to HY-160799 concentrations previously reported, ranging from 1-10 µM (Supplemental Fig. S3 A-B) [22]. For treatment, *Sdhb^+/+^* and *Sdhb^-/-^* imCCs were plated on 6-well plates at 50% confluence and allowed to settle overnight. The next day, normal growth media were switched with 2 mL of fresh treatment media at 1.5, 3, 5, 7 and 10 µM drug concentration for MLN4924 (NEDD8 inhibitor) and 1, 3, 5 and 10 µM for HY-160799 (UBE2F inhibitor). Each concentration was tested in triplicate in the presence of DMSO controls. Fresh drug-containing media were provided daily for the duration of the treatment period.

### Measuring cell proliferation

Proliferation for shRNA studies was reported over a 7-d growth period for each line after the 10^th^ cell doubling when sufficient UBE2M and UBE2F reduction was observed. Proliferation for chemical inhibition studies was determined over a treatment period of two doublings, i.e. 2 d for *Sdhb^+/+^* cells and 8 d for *Sdhb^-/-^*cells [13]. The results of this method did not vary from the pilot experiment, in which proliferation was established over 7-d treatment period. Results were relatively consistent when 2 doublings were accounted for versus when cells were treated for 7 consecutive days regardless of doubling time (Fig. 3B-C and Supplemental Fig. S4). Cell counting was performed label-free, where N_0_ represents initial cell count, *N_t_* is the final count and *T* is the number of days.

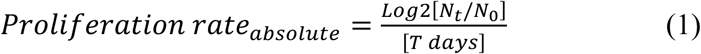

Relative growth rate was determined by dividing absolute growth rate of treated cells over that of untreated cells.

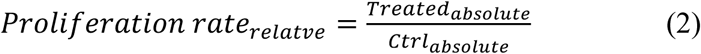

### Western blotting

A standard western blotting protocol was followed to assess various protein levels throughout the screening and validation processes. In all cases, cells were trypsin-harvested at 80% confluence from T-175 flasks (approximately 3 million cells), washed in PBS and lysed in 150 μL cold RIPA buffer containing protease and phosphatase inhibitor cocktail (Santa Cruz Biotechnology # sc-24948). Samples were incubated on ice for 30 min with gentle vortex mixing every 10 min then subjected to centrifugation at 15,000×g for 15 min at 4 °C. After transferring the supernatant containing protein to new 1.5-mL tubes, protein quantification was carried out using a BCA protein assay kit (Pierce # A55864) according to manufacturer’s protocol. Samples were heated to 70 °C for 10 min after the addition of reducing agent and an appropriate volume of 4× LDS denaturation buffer. Denatured samples (30-50 µg total protein) were subjected to electrophoresis through NuPAGE 10% Bis-Tris protein gels in MES-SDS running buffer (Invitrogen #NP0060) at 150 V for 1 h. Membrane transfer was performed using PVDF membranes (BIO-RAD #1620174) according to the Novex Western Transfer Apparatus protocol in NuPage transfer buffer (Invitrogen #NP00061) containing 20% methanol. at 4 °C (30 V, 245 mA) for 90 min. Membranes were then blocked with 3% non-fat milk for 1 h at room temperature, followed by washing in TBST buffer. All blots were run simultaneously using the same protein extracts and master-mixes and assigned antibody randomly if size or cross-reactivity were expected to interfere with co-staining. Primary antibody dilution buffer was made by combining 7.5 mL TBST, 2.5 mL 4% BSA, and 250 μL 0.5% sodium azide with appropriate antibody. After 24-48h of incubation at 4 °C, blots were washed three times in TBST before staining with IRDye^®^ 680rd goat anti-rabbit IgG (LI-COR #926-68071, 1:15000) or 800cw goat anti-mouse IgG secondary antibody (LI-COR #926-32210, 1:15000) antibodies in TBST with 3% non-fat milk for 1h at room temperature prior to imaging. Blots were imaged with their appropriate settings on an Amersham Typhoon™ 5 biomolecular imager (Amersham Typhoon, Uppsala, Sweden).

Primary antibodies used were: α-CRISPR-Cas9 (Abcam #ab210571; 1:1000); α-SDHB (Abcam #459230; 1:2000), α-GAPDH (Abcam #ab8245; 1:5000), α-UBE2M (Abcam #ab109507; 1:5000), α-UBE2F (Proteintech #17056-1-AP; 1:5000), α-CUL5 (Abcam #ab184177; 1.5:5000), α-NEDD8 (Invitrogen #PA5-17476; 1.5:5000) and α-NOXA (Boster bio #M02287; 1:1000).

### Data analysis

CaRpool analysis was performed using the method of *Winter et al., 2016*, mapped to CP0044 (sgRNA) Brie Genome 1-4 referencing GRCm38 GRC from The Broad Institute’s Genetic Perturbation Platform [41]. MaGecK was performed according to previously published work [42]. In short, the algorithm ranks all sgRNAs per gene (encompassing guides for any transcriptional start sites) using p-values derived from a mean variance model. To ascertain which genes are outstanding in enrichment or depletion, a robust ranking aggregation (RRA) algorithm is then employed, leveraging p-value and false discovery rate (FDR) statistics. Guides that scored less than 100 initial folds of coverage were excluded from the analysis.

DepMap effect scores were extracted from (https://depmap.org/, DepMap Public 25Q2 version) and selected for kidney renal cell carcinoma cell lines (n=57). Average effects scores were then calculated for each individual gene across selected cell lines and tested for their association with MAGeCK-RRA CRISPR SDHB/C screen scores.

Bar graphs and relevant statistical significance of different groups were generated and analyzed using GraphPad Prism7 (GraphPad Software, San Diego, CA, USA). Specifics for each graph, including tests and significance are reported as part of the figure legends.

## Abbreviations

caRpools: analysis pipeline for pooled CRISPR screens
CRISPR: clustered regularly interspaced short palindromic repeats
DepMap: dependency map
FDR: false discovery rate
GRC: genome reference consortium
imCCs: immortalized mouse chromaffin cells
iMEFs: immortalized mouse embryonic fibroblasts
LOH: loss of heterozygosity
MAGeCK: model-based analysis of genome-wide CRISPR-Cas9 knockouts
MEFs: mouse embryonic fibroblasts
NT: none-targeted
PCR: polymerase chain reaction
PGL: paraganglioma
PHEO: pheochromocytoma
PPGL: pheochromocytoma and paraganglioma
PRRT: peptide receptor radionuclide therapy
RRA: robust rank aggregation
SDH: succinate dehydrogenase
shRNA: short hairpin RNA

## Acknowledgments

The authors acknowledge the consultatory and technical assistance of the staff at the Mayo Clinic Division of Laboratory Genetics and Genomics Laboratory. We wholeheartedly thank Matthew Schellenberg Ph.D. and Scott H. Kaufmann, M.D., Ph.D. of Mayo Clinic for sharing reagents and for crucial discussions. Members of the Maher laboratory are also acknowledged. BioRender.com was used to assist in the generation of some figures.

## Data and resource availability

Materials and data are available upon request to the authors. CRISPR screen data deposition pending.

## Author contributions

FJAK, MJE, JRH and LJM conceived and designed the experiments; FJAK carried the experiments; FJAK, MJE and CMdeAC analyzed data; FJAK and LJM wrote the manuscript; JF provided research materials and manuscript editing.

