## Supplemental material for "Unbiased CRISPR Synthetic Lethal Screening for Genetic Vulnerabilities in Succinate Dehydrogenase (SDH)-loss Model of Paraganglioma"

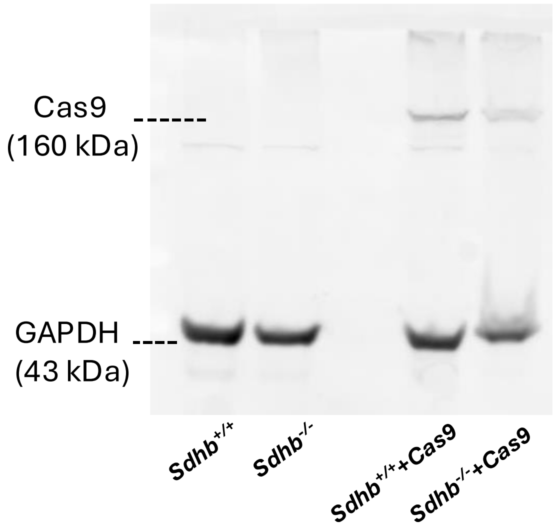


**Supplemental Figure S1:** *Immunoblot assessment of Cas9 expression in Cas9-transduced cells compared to their parental lines.* Western blot was used for detection of  Cas9 (160 kDa) in cell lysates of parental and Cas9-transduced imCC lines. GAPDH (43 kDa) was used s loading control.

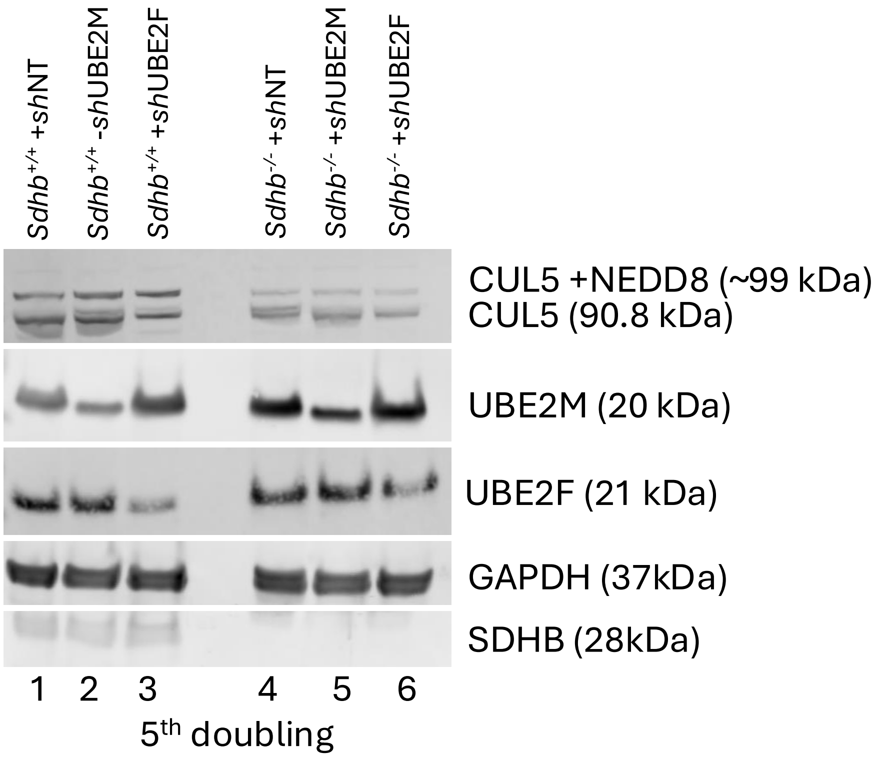


**Supplemental Figure S2.** *Western blot analysis of protein expression in Sdhb$^{+/+}$ and Sdhb$^{-/-}$ cells 5 doublings after shRNA knockdown treatment***.** Western blot assessing the expression of several proteins in *Sdhb*$^{+/+}$ and *Sdhb*$^{-/-}$ cells, each treated with non-targeting shRNA (+shNT), shRNA targeting UBE2M (+shUBE2M), or shRNA targeting UBE2F (+shUBE2F). The proteins detected include CUL5 (90.8 -99 kDa), UBE2M (20 kDa), UBE2F (21 kDa), GAPDH (37 kDa), and SDHB (28 kDa). GAPDH serves as a loading control, and SDHB confirms the *Sdhb* status. Samples were collected after the fifth cell doubling for each line, when moderate change in targeted protein expression was observed compared to after the tenth doubling (Fig. 2D).

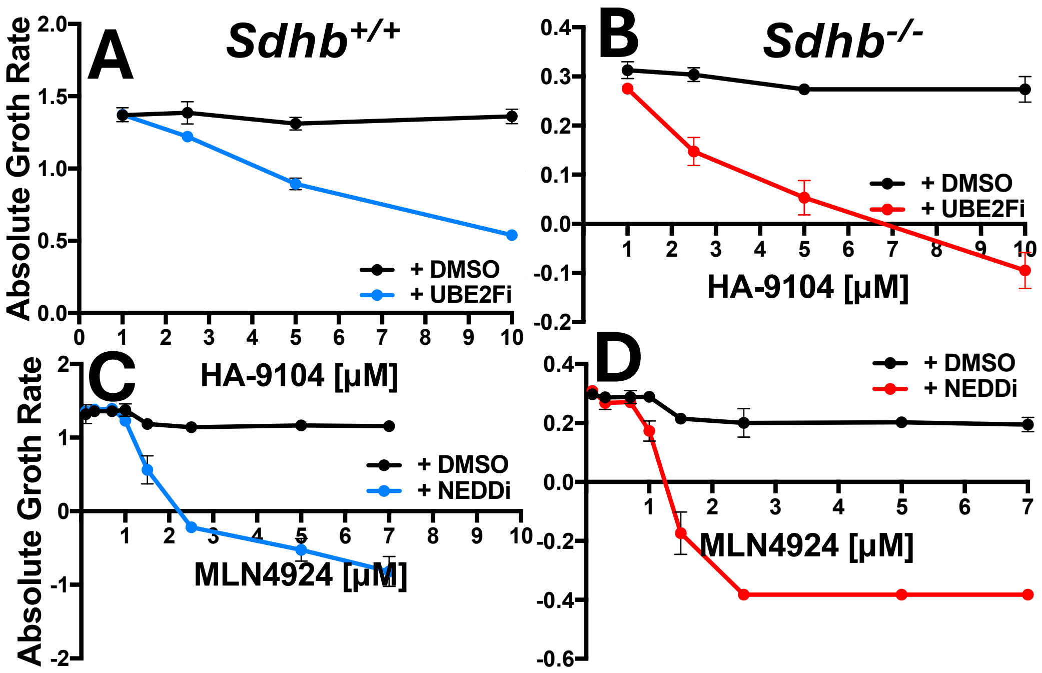


**Supplemental Figure S3.** *Absolute growth rate for data reported in Fig. 3B-C.* Raw data were calculated over two cell doublings (2 d for *Sdhb*$^{+/+}$ [A and C], 8 d for *Sdhb*$^{-/-}$ cells [B and D]) using equation 1 reported in methods. Negative absolute growth rate signifies a decrease in cell number over the measured period. *Sdhb*$^{-/-}$ cell data: red lines; *Sdhb*$^{+/+}$ data: blue lines; black lines represent a matching concentration of DMSO solvent.

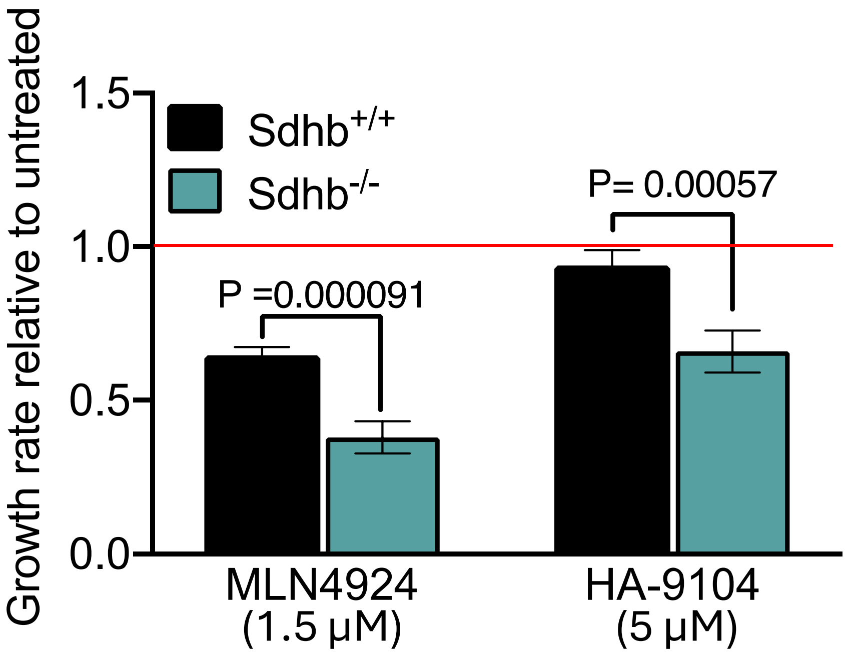


**Supplemental Figure S4.** *Effects of 7-d treatment with anti-neddylation drugs at the indicated concentrations***.** Shown is growth rate of *Sdhb*$^{+/+}$ (black bars) and *Sdhb*$^{-/-}$ (teal bars) cells relative to DMSO controls. Cells were treated with either MLN4924 (1.5 µM) or HA-9104 (5 µM) for a period of 7 d for each line. Red line indicates a relative growth rate of 1.0, representing growth of untreated cells. Error bars represent the standard error of the mean after error propagation.
